# A DNA base-specific sequence interposed between CRX and NRL contributes to *RHODOPSIN* expression

**DOI:** 10.1101/2023.12.22.573078

**Authors:** Rosa Maritato, Alessia Medugno, Emanuela D’Andretta, Giulia De Riso, Mariangela Lupo, Salvatore Botta, Elena Marrocco, Mario Renda, Martina Sofia, Claudio Mussolino, Maria Laura Bacci, Enrico Maria Surace

## Abstract

The binding affinity of transcription factors (TFs) for their cognate DNA sequences controls gene expression. DNA determines the recruitment and positioning of TFs; whether it plays other roles is unknown. Here we found that the specific 22-bp sequence interposed between the CRX and NRL TFs in the proximal promoter of *RHODOPSIN* (*RHO*) largely controls the expression levels of *RHO*. Mutagenesis of this DNA-linker sequence resulted in wide variation in gene expression. In contrast, reciprocal exchange of human and murine *RHO* elements conferred species-specific expression levels. Targeting the DNA-linker with equal orthogonal DNA-binding proteins activates or represses *RHO* expression depending on its orientation relative to CRX and NRL binding sites. We conclude that DNA itself adds to TF activity through a code that determines optimal levels of *RHO* expression.

## Introduction

DNA plays a dual role in gene expression. It encodes for regulators of transcription such as transcription factors (TFs) and allows their positioning in the transcriptional machinery by bearing specific transcription factor binding sites (TFBSs) (*1,2*). The fundamental interaction that generates DNA-protein binding specificity and eventually gene expression, is generated by the specific affinity of TF protein contacts with specific DNA bases within TFBSs (*3-5*). However, in metazoans, the need for tight control of the level, pattern, and timing of gene expression in different cell types has meant that interactions between TFs and their cognate TFBSs have evolved in a more complex and diverse manner (*6*). One TF binds multiple genes, and one gene is regulated by multiple TFBSs, and thus by multiple TFs-TFBS interactions (*7*). TFBSs contain DNA sequence variation that exceeds that present in the evolutionarily more constrained coding sequence of TFs, in a model of co-evolution in which TFBSs evolve faster than TFs (*8-10*). TFBSs are short degenerate DNA sequences (also known as motifs) that occur as monomeric (*11,12*), dimeric, and overlapping sequences embedded in cis-regulatory elements, so that the DNA context surrounding them can also influence how they are bound by TFs (*13-17*). On the other hand, TFs couple TF-TF interactions (TF-TFs) to cooperatively and combinatorially (*18,19*) recognize and bind TFBSs through protein-(*18, 20*) or DNA-protein-driven mechanisms (*13, 21-24*). On a larger scale, TFs and TFBSs combine their binding modes functionally to determine gene expression. The number, position, helical DNA face and binding site (strand) of TFBSs and their flanks are recognized by multiple TFs that can bind independently (*11,12,25*), or cooperatively by co-binding (*26-28*), resulting in qualitative and quantitative binding at the promoters and enhancers (*29-31*). These different modes originate and resolve in one recognized basic mechanism, TF-DNA binding affinity (*32,33*). The dynamic interactions that regulate DNA-protein recognition passing through chromatin status and interaction of TFs with nucleosomes (*34,35*), eventually lead to rates of association and dissociation of DNA-binding proteins to and from DNA, which in turn determine cell-specific transcriptional activity (*36-38*). Therefore, the high to low affinity ranges of DNA-protein interactions (the number and types of TFBSs and TF concentrations and types (*39-41*)) account of gene expression from noise (*42*) to gene expression programs (*43,44*). However, it is unknown whether evolution has generated an additional level of DNA-protein interaction in gene regulation.

The human *RHODOPSIN (RHO*) gene encodes for a G-protein-coupled receptor expressed exclusively in retinal rod photoreceptors and located in their outer segment (OS) (*45*). RHO in the OS intercepts and converts light photons into electrical signals through the phototransduction cascade and, when mutated, causes retinitis pigmentosa (RP) and other blinding disorders (*46, 47*). *RHO* expression during development increases linearly and after birth becomes highly and constantly transcribed in rod photoreceptors (*48*). The *Rho* gene is also tightly regulated, as both low and high expression levels can lead to photoreceptor degeneration (*49,50*). Despite the presence of two conserved enhancers (*Rho Enhancer Region* (*RER*) (*51*) and *CRX-Bound Region 1*(*CBR*) (*52*)), *Rho* expression in the adult retina is mainly controlled by the adult-specific accessible regions at the proximal promoter (*53, 54*).

We previously have shown that, both a synthetic zinc finger protein (ZF-DB) and an ectopically expressed transcription factor (KLF15) block *Rhodopsin* (*Rho*) expression when delivered *in vivo* to rod photoreceptors of adult porcine retina by Adeno-associated virus (AAV) vectors (*55,56*). ZF-DB and KLF15 bind 20- and 9-bp DNA sequences, partially overlapping (6 bp), between two key TFs for *RHO* expression, the Cone Rod Homeobox protein (CRX) (*57*) and the Neural Retina-specific Leucine zipper protein (NRL) (*58*) in the proximal promoter region of *RHO* (-82 to -62 and -84 to -75 from the transcription start site, TSS, respectively; (Fig.1A)). ZF-DB and KLF15 repression of *Rho* caught our attention for two main aspects: (i) ZF-DB and KLF15 bind sensitively a DNase-unprotected (*59,60*) region at 20-fold lower concentrations than those of CRX and NRL, and selectively, as low number of transcriptional off targets are induced (*55,56*); (ii) ZF-DB and KLF15 resulted in *Rho* repression despite their biochemical differences, ZF-DB possesses exclusively DNA-binding properties (without effector domains), KLF15 is a fully competent TF (*61*), overall suggesting that a very compact DNA regulatory sequence near to TSS controls *Rho* expression in terminally differentiated rods. Thus, ZF-DB and KLF15 within *Rho* native chromosomal context *in vivo*, upon binding to an accessible DNA sequence between CRX and NRL are sufficient to overcome the transactivation effects of CRX and NRL and in turn block *Rho* expression. These observations led us to hypothesize that CRX and NRL TFs pair additionally to binding, interact with the specific DNA sequence interposed between them by a novel mechanism, which ultimately contributes to determining *Rho* expression levels.

## RESULTS

### Forty-nine bp containing CRX and NRL BSs (RHO regulatory unit, RRU) control Rho expression in adult retina

ZF-DB and KLF15 target a sequence between two extensively validated regulatory elements, BAT-1 and NRE (*52, 59, 62*), which are bound by the retinal-specific TF, CRX (*57*) and by the rod photoreceptor-specific NRL (*58*), respectively. CRX-NRL pair operate synergistically to transactivate numerous rod-specific genes, including *RHO* (*52,62*). On this basis, we assessed whether the 49 bp long regulatory elements encompassing the CRX and NRL binding sites and the interposed linker sequence bound by ZF-DB and KLF15 may represent a regulatory unit (here named *RHO* regulatory unit, RRU) controlling *RHO* expression (Fig. 1A). To this end, we generated constructs for an *in vivo* AAV reporter assay (Fig. 1A). This *in vivo* reporter assay (*52*) enables to link variation of a DNA regulatory sequence to the activity of a reporter gene, here, the enhanced *Green Fluorescent Protein* reporter gene (*eGFP*) by measuring gene expression variation output derived by an input sequence delivered by AAV to the target tissue, here the adult mouse retina (Fig. 1A). We thus, generated a human *RHO* proximal promoter fragment of 259 bp (hRHOp: -164 bp from TSS and 95 bp of 5’UTR), containing the 49 bp of the RRU, to drive the expression of the *eGFP* (Fig. 1A). Fifteen days after subretinal delivery of AAV-hRHOp-eGFP, gene expression levels measured by RT-PCR, were shown evident and immunofluorescence analysis showed *eGFP* gene expression confined to rod photoreceptors. To dissect the contribution of DNA sequence determinants of RRU, we first mutated the CRX and NRL binding sites. AAV-hRHOp-eGFP carrying individual mutations in mCRX1, mCRX2 or NRL binding sites showed that each binding site is required to sustained *eGFP* reporter expression (Fig. 1B). Next to assess whether RRU is necessary for *RHO* expression, we either delete (Prom-Q) or duplicated (Prom-S) the RRU (Fig. 1C). AAV-Prom-Q abolished reporter expression levels, whereas AAV-Prom-S duplication almost doubled *eGFP* expression. These results suggest that RRU (a 49 bp long *RHO* compact regulatory element containing CRX and NRL binding sites and the interposed linker sequence) contains key features necessary to control both levels and expression pattern restricted to rod-photoreceptors expression in adult mice.

**Fig. 1.**
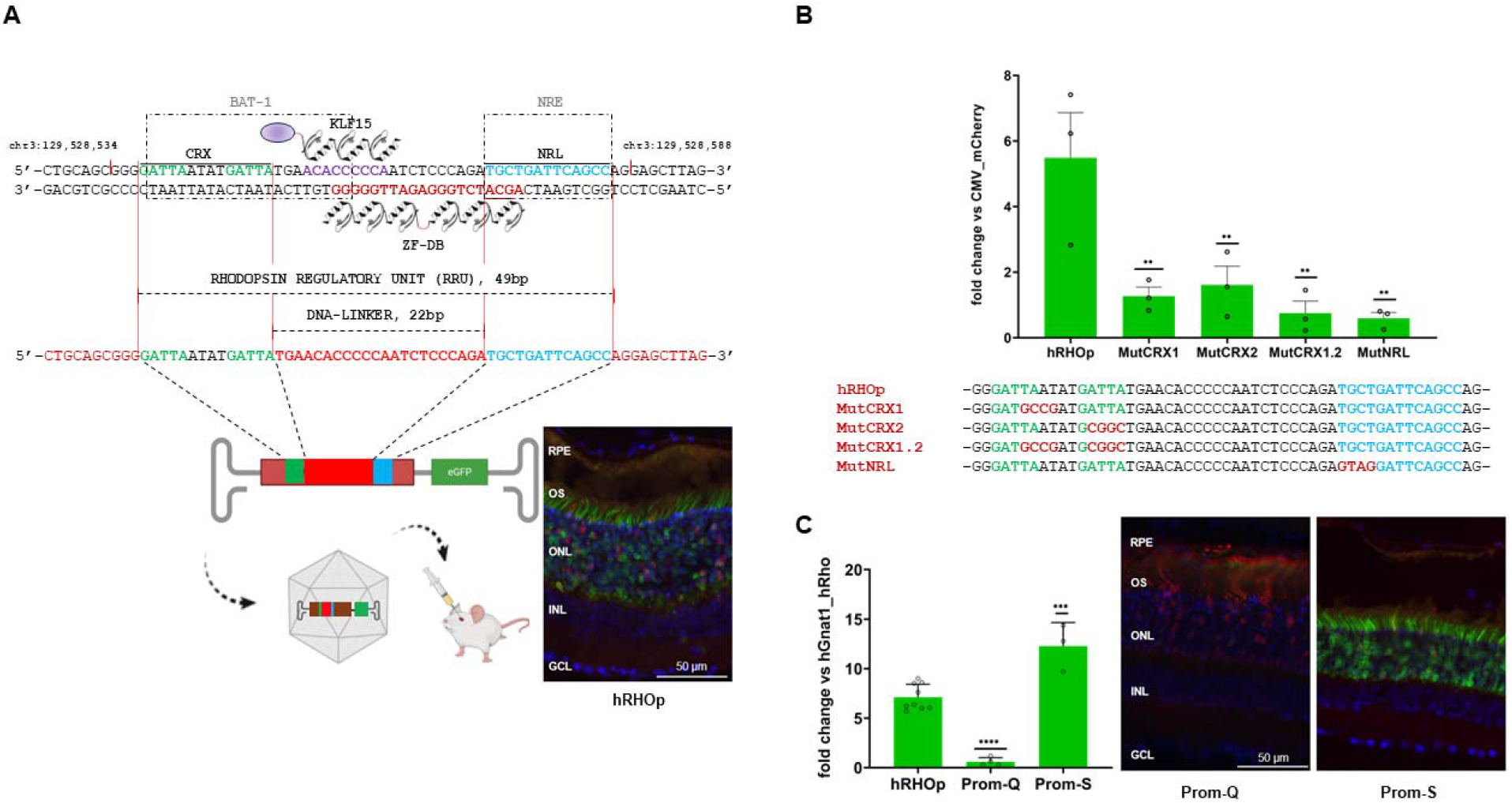
Forty-nine bp containing CRX and NRL BSs control *RHO* expression in adult retina. (**A**) The proximal promoter region of the human *RHO* containing the Rho Regulatory Unit (hRRU, chr3:129,528,537-129,528,585). It is composed of the BAT-1 and NRE elements (dashed grey boxes), CRX and NRL TFs binding sites (in green and light blue, respectively) and DNA-Linker (in red). The “footprints” onto hRHOp of the synthetic zinc finger protein (drawing of the six zinc-finger array of ZF-DB below the negative strand, amaranth) and the TF KLF15 (drawing of KLF15 including the effector domain in violet above the positive strand). Below, schematic representation of gene reporter assay with of Adeno-associated virus (AAV) vector carrying *eGFP* under the control of the *RHO* proximal promoter (hRHOp) injected sub-retina in adult mice, on the right-handed panel immunofluorescence analysis of hRHOp 15 days post-injection. (**B**) Histogram, quantitative PCR analysis (qPCR), *eGFP* expression levels 15 days after AAV-eGFP *in vivo* retinal delivery in adult mice (methods). (**C**) qPCR upon removal of hRRU (Prom-Q) and doubling the hRRU (Prom-S), and, immunofluorescence analysis on Prom-Q, and Prom-S co-injected with a AAV vector encoding for mCherry. RPE, retinal pigment epithelium; OS, outer segment; ONL, outer nuclear layer; INL, inner nuclear layer; GCL, ganglion cell layer.

### Changes of the DNA sequence interposed between CRX and NRL leads to differential reporter gene expression

To dissect the DNA sequence determinants within RRU, we restricted the analysis to the 22 bp long DNA sequence intervening between CRX and NRL, from now on termed the DNA-linker (Fig. 1A), and its relationship with the flanking CRX and NRL binding sites. To do so, we generated constructs carrying changes in CRX and NRL binding sites order, orientation, and the spacing between them (the length of the DNA-linker), referred as the regulatory syntax (*37*) and tested them *in vivo* with AAV reporter assay. Mutual changes of CRX and NRL binding sites order (AAV-Prom G-1), and concurrent change of order and orientation (AAV-Prom G-2) resulted in less than half, and abolished reporter expression levels, respectively, relative to control (Fig. 2A). To manipulate the spacing between CRX and NRL binding sites, we generated three distinct non overlapping deletions of 5 bp, Del-5’, Del-C and Del-3’, in their respective AAV-reporter vectors, within CRX and NRL DNA-linker. These deletions entail a half DNA helical turn (10.5 bp represents an entire DNA turn) of the CRX and NRL binding sites, arranging them on the opposite sides relative to native configuration. *In vivo* reporter gene expression varied in these three constructs from null levels (AAV-Del-3’) to levels equivalent (AAV-Del-5’) to the native sequence (Fig. 2A). The fact that the same reduction of spacing resulted in differential *eGFP* expression levels, with an apparent decrease in the 5’ to 3’ direction of the DNA-linker, points that the source of the variation is contained into CRX and NRL linking sequence, thus, we next mutated its DNA base sequence. We generated both contiguous (7 to 10 bp) and a single base substitution. These changes resulted in diversified levels of reporter gene expression (Fig. 2C). Compared with deletions, in this case only the first 5’ mutagenesis of AAV-Prom-N retains some of the activity, while a few bases away from the 3’ inner flank of the DNA-linker (AAV-Prom-Me and AAV-Prom-3a) reporter expression begins to be greatly reduced. Thus, deletions and base substitutions within the DNA-linker lead to highly differentiated gene expression (Fig. 2C and Fig. 2D). This suggest that DNA base content and DNA-linker length are key features for the function of RRU. However, these effects may be due to the generation of *de novo* binding sites for either activators or repressors. To rule out this possibility we scanned each sequence for the respective BSs from the HOCOMOCO human collection of TFs (see Methods) (*63*). The NRL and CRX BSs were consistently retained in the flanks of the DNA-linker in all hRHOp artificial promoter sequences tested, thus, compatible with previously described occupancy on the human *RHO* promoter (*60*). On the contrary, in the DNA-linkers of both wild-type and synthetic promoters we were not able to identify any match with TFs expressed in the human retina (Table S2). To further rule out the possibility of generating novel BSs we inverted the 5’ and 3’ orientation (Prom-L; Fig. 2B). This arrangement affects only the 3’ and the 5’ flanks of CRX and NRL binding sites, respectively (Fig. 2B), while ensuring both the conservation of the DNA-linker sequence and its plus-minus DNA strands configuration relative to CRX and NRL BSs. Notably, in this arrangement the reporter expression levels were strongly reduced (Fig. 2B). Finally, immunofluorescence analysis showed that these changes of levels of reporter expression across different mutants corresponded to a variation of the intensity of eGFP expression and maintenance of the same eGFP pattern, restricted to rod-photoreceptors (Fig. 1A, C and Fig. 2D). Collectively, these results suggest that without the apparent intervening action of other retina-specific TFs, the DNA linker connecting CRX and NRL controls the levels but not the pattern of *RHO* expression. Thus, the source of variation in *RHO* reporter expression depends on the syntax of CRX and NRL BSs (order, orientation, and distance of TFBSs), and in addition on the base composition of the DNA-linker sequence interposed between CRX and NRL (Prom-L).

**Fig. 2.**
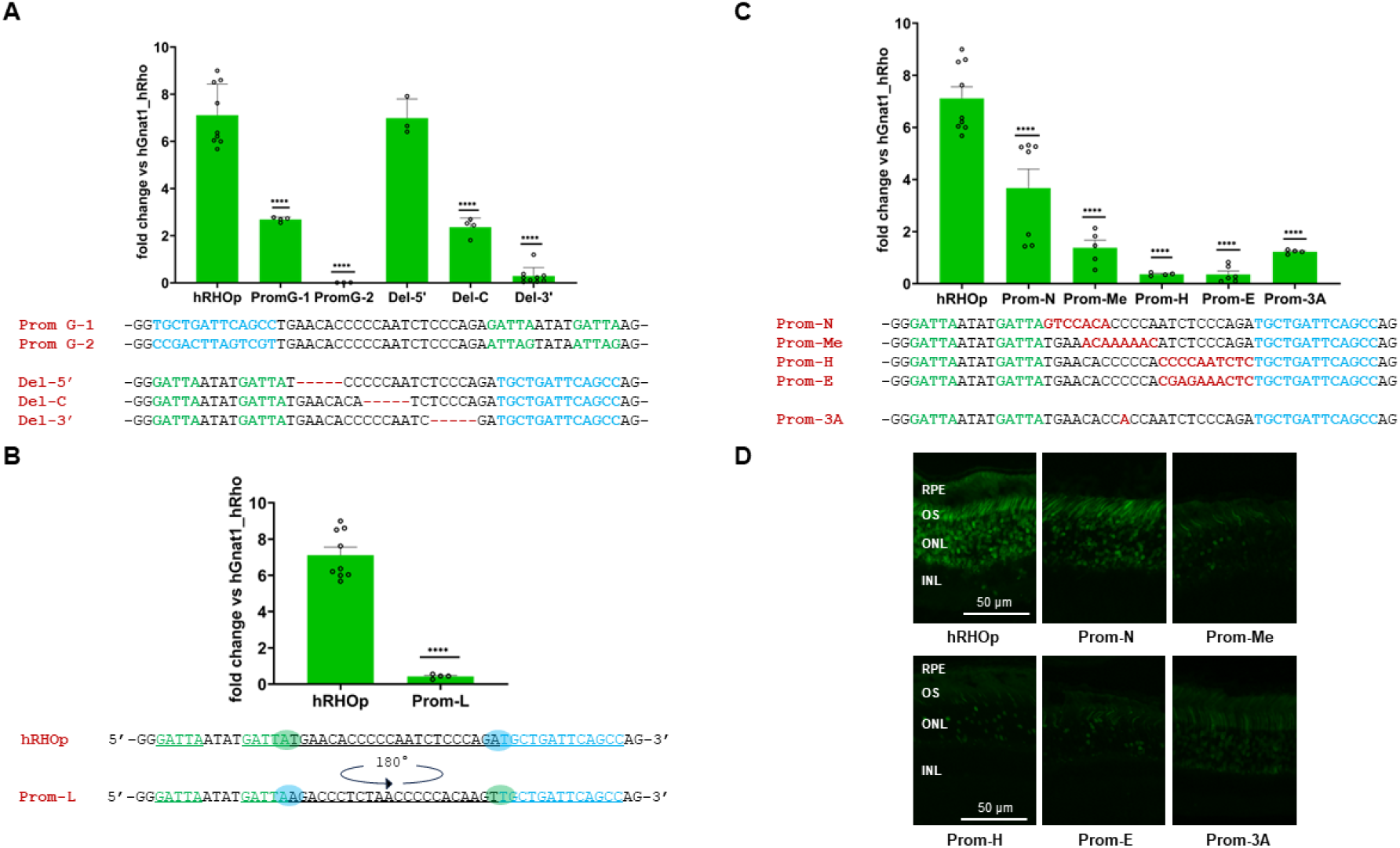
Changes of the DNA sequence interposed between CRX and NRL lead to uncorrelated reporter gene expression. **(A)** Sequences, and histogram, qPCR analysis, *eGFP* expression levels 15 days after AAV-eGFP in vivo retinal delivery in adult mice (methods), showing the impact of change of the order, orientation, and spacing, of CRX and NRL BSs. (**B**) Quantitative effects (qPCR) of DNA-linker sequence cloned in the 5’ and 3’ orientation (sequence, AAV-Prom-L). (**C**) Sequences, and qPCR of nucleotide substitutions in the DNA-Linker. (**D**) Immunofluorescence analysis on AAV-hRHOp, AAV-Prom-N, AAV-Prom-Me, AAV-Prom-H, AAV-Prom-E and AAV-Prom-3A. RPE, retinal pigment epithelium; OS, outer segment; ONL, outer nuclear layer; INL, inner nuclear layer; GCL, ganglion cell layer.

### Human and murine RRUs diverge but lead to correlated reporter gene expression

Next, we compared the DNA sequence features of RRU along the mammalian phylogeny. The alignment of 100 mammalian genomes shows that, apart from rodents, the two CRX and the NRL binding sites are conserved (Fig. 3A, B and Table S2). Another noteworthy feature is the conserved spacing of 22 nucleotides of the DNA-linker. However, alternative nucleotide changes are present in several mammals, although not completely evenly distributed (the central sequence is more conserved than the 5’ and 3’ regions), including the internal bases flanking core CRX and NRL binding sites (Fig. 3A). Therefore, given conserved spacing, nucleotide sequence varies across mammalian phylogeny (Fig. S3). Unlike other mammals, in rodents, and particularly in mouse and rat, RRU has one CRX binding site, and the DNA-linker distance has one more nucleotide (23 bp), while nucleotide sequence of the DNA-linker diverges extensively (Fig. 3A). Thus, we compared the human and murine homolog sequence. To study murine RRU (mRRU), we generated AAV reporter constructs with the proximal murine rhodopsin promoter (mRhop, -164 bp from the TSS and -78 bp from the 5’UTR). The murine promoter resulted in slightly lower reporter gene expression than the human counterpart (Fig. 3C). However, reciprocal replacement of human RRU with murine RRU in human *RHO*-promoter (hRHOp) and of murine RRU with human RRU in mouse-*Rho* promoter (mRhop) restored the species-specific reporter expression levels. This suggests that despite the divergences between the DNA-linker, human and murine RRUs are functionally equivalent and relatively independent from both the embedded motif and the background sequences (*17*). Furthermore, between humans and mice, nucleotide divergence in the DNA-linker corresponded to similar levels of reporter expression, supporting that, unlike synthetic constructs, a functional correlation may be obtained via an evolution-driven nucleotide arrangement. To further explore and isolate the properties of the human and murine DNA-linkers we swapped the 5 bases in the middle of them. These sequences are highly divergent, GC-rich in human and AT-rich in mouse (Fig.3D). In addition, these bases are centered in the DNase-unprotected sequence (59, 60). AAV constructs containing the human-*RHO* promoter carrying the mouse 5 bp element (ATGAT, AAV-hRHOp-mAT) and the mouse-*Rho* promoter carrying the human 5 bp element (CCCCA, AAV-mRhop-hCG), showed a precise inversion of the expression levels, the human *RHO*-promoter with the mouse sequence (hRHOp-mAT) equals mouse-*Rho* promoter levels, whereas, the murine *Rho*-promoter (mRhop-hCG) equals the human *RHO*-promoter (Fig.). This suggests that an even shorter sequence within the DNA-linkers is responsible for the species-specific *Rho* expression levels.

**Fig. 3.**
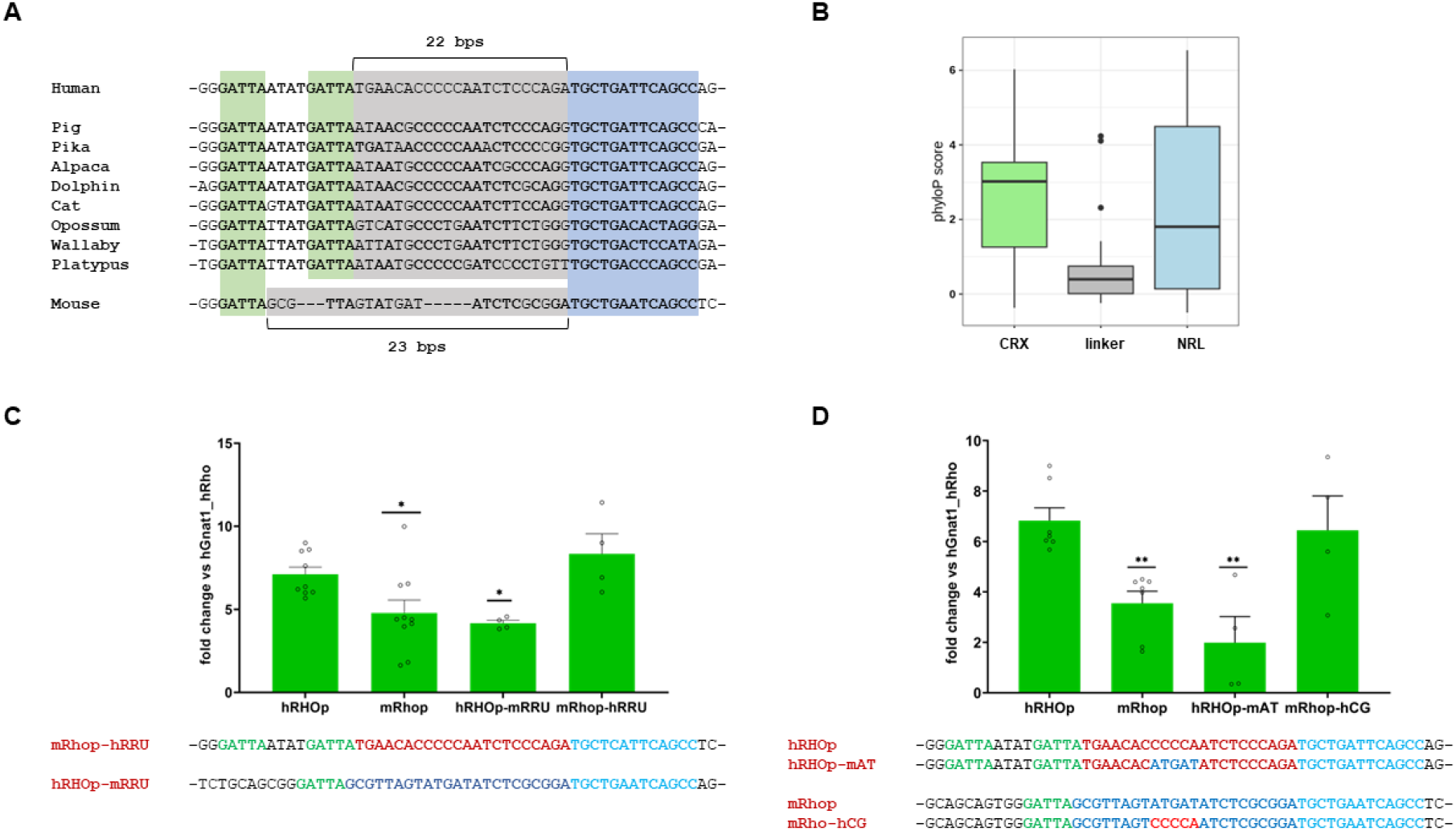
Human and murine DNA-linkers diverge but lead to specie-specific *RHO* reporter expression. **(A)** Multiple sequence alignment for representative mammalian species from UCSC track. The CRX and NRL binding sites are highlighted in green and blue, respectively. The DNA linker is highlighted in grey. Conserved bases are indicated in bold. (**B**) Distribution of PhyloP score of DNA bases in the NRL (green box) and CRX binding sites (blue box) and in the DNA-linker of the human RRU. PhyloP score indicates the degree of conservation of a DNA base in vertebrates. (**C**) Sequences and qPCR analysis of mutual changes of hRRU (AAV-mRhop-hRRU) and mRRU (AAV-hRHOp-mRRU) insertion in murine and human *RHO* promoters, respectively. (**D**) Sequence and qPCR analysis of the human *RHO* promoter with 5 bp of central murine DNA-linker (ATGAT, AAV-hRHOp-mAT) and mouse *Rho* promoter with 5 bp of central human DNA-linker (CCCCA, AAV-mRhop-hCG).

### Interference with DNA-binding proteins results in effects dictated by the DNA-linker orientation

To determine whether these cis-regulatory sequence properties are retained, in isolation, in an unrelated trans-regulatory system in which retina-specific genes are not expressed, including CRX and NRL, we co-transfected HEK293 cell line with the same plasmids used to generate AAV vectors and those encoding for CRX and NRL (Methods). hRHOp showed synergistic activation upon co-transfection with CRX and NRL and similar gene expression variation as observed *in vivo* on several constructs (Fig. S1B). Next, we interfered the DNA-linker activity with the DNA-linker binding proteins which we have previously used to repress RHO expression *in vivo* (*55, 56*).

Since the first of the six fingers of the ZF-DB protein intrudes into the NRL-binding site (Fig. 1A) we removed it obtaining ZF-DB-5 (Fig. 4B). This ZF-DB-5 mutant, which has shown similar *RHO* repression properties as that of the ZF-DB, in RHO-P347S mice retina *in vivo* (Fig. S2) (*55*), when transfected with hRHOp *in vitro*, repressed *RHO* reporter expression. Next, to get insights into the potential mechanisms underlying the action of the DNA-linker, we tested whether the DNA-linker L (Prom-L) which carries the DNA-linker sequence cloned in the 3’-5’ direction, thus maintains the same binding sites for ZF-DB-5 and KLF15 (Fig. 4A and B). Surprisingly, Prom-L, which has shown to strongly reduce *RHO* reporter activity when co-transfected with ZF-DB-5 and KLF15 promoted its transactivation (Fig. 4A and B). Notably, ZF-DB-5 without a transactivation domain increased the expression of the reporter more than KLF15, which possesses catalytic domain. These results suggest that interfering the DNA-linker sequence with orthogonal proteins results in down- or up-regulation of gene expression, depending on the orientation of the DNA-linker sequence between CRX and NRL binding sites.

**Fig. 4.**
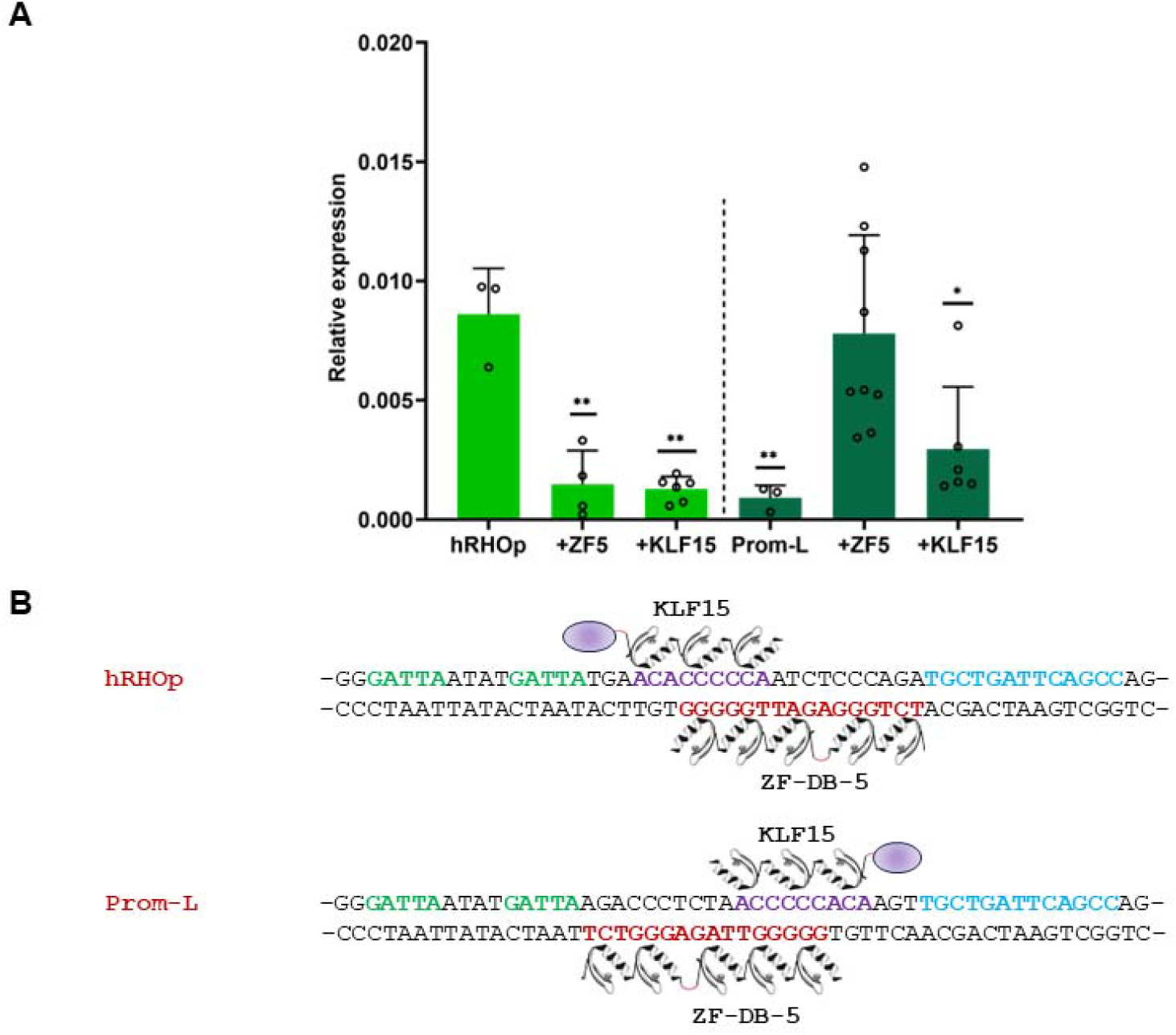
Interference with DNA-binding proteins results in effects dictated by the DNA-linker orientation. **(A)** Quantitative effects (qPCR) of transfecting ZF-DB-5 or KLF15 in HEK-293 cell line with hRHOp or Prom-L promoters. (**B**) ZF-DB-5 and KLF15 drawing bound to the hRHOp or Prom-L promoters sequences.

## DISCUSSION

We found that a 49-bp-long sequence of the human *RHO* proximal promoter, containing binding sites (BSs) for CRX and NRL, is a functional discrete cis-regulatory unit (RHO regulatory unit, RRU) that controls *RHO* expression levels in the adult rod-photoreceptors of the retina. This result is in accordance with previous observations which have shown that short proximal sequences of *RHO* promoter enable *Rho* expression *in vivo* in adult mouse rod-photoreceptors (*53, 54, 64*) and that *Rho* enhancers are largely unnecessary. These results are also consistent with the effect we showed *in vivo*, in native chromosomal context, with two transcriptional repressors, ZF-DB or KLF15 (*55, 56*). These DNA-binding proteins when expressed by AAV in rod-photoreceptors, independently, bind a partially overlapping 23 bp sequence at the *RHO* proximal promoter, blocking *RHO* expression. This suggests that, in terminally differentiated adult photoreceptors, the source of *RHO* expression levels is controlled within an even shorter sequence contained in the RRU of the proximal promoter of *RHO*, the “DNA-linker”. In fact, we found that within RRU, the base specific composition and length (22 bp) of the DNA-linker, the sequence interposed between CRX and NRL of the human *RHO* promoter, controls *RHO* expression.

This allowed us to intercept a potential additional property of DNA-protein interaction. We first found that the shortening (5 bp) of three different consecutive regions of the DNA-linker, which leads to a different DNA sequence between the CRX and NRL binding sites (Fig. 2A), caused a differential *eGFP* expression in the retina, *in vivo* (Fig. S1A). Similarly, mutagenesis of the DNA-linker resulted in highly differentiated *eGFP* expression, consistent with that observed by others in the homologous sequence of the murine *Rho* promoter (*65*). Notably, the sequence derived by both these shortened and mutagenized DNA-linkers did not generate novel binding sites for TFs expressed in the human retina. These results were also confirmed by immunofluorescence analysis, which showed that the expression of eGFP was found invariably restricted to photoreceptors, as the native sequence, albeit with different intensities, thus suggesting that the mutated linkers do not allow binding of additional TFs. Complementarily, when guided by evolution, we found that the relative optimal expression levels of the human and murine RHO proximal promoters were determined by the specific composition and length of the DNA sequence of their respective DNA-linkers. Human and murine RHO proximal promoters show a slightly different reporter expression levels in favor of the human promoter, in our experimental setting. Notably, swapping their respective RRUs, translated functionally in an inversion of promoter strengths. Furthermore, we showed that the exchange of the 5 bp divergent sequences between the human and mouse, in the middle of their DNA-linkers, enables an exact species-specific segregation of their respective *RHO* promoter strength (Fig. 3D). Therefore, in contrast to shortened and mutagenized DNA-linkers, these naturally occurring divergent human and murine DNA-linkers promoted RHO gene expression in a correlated manner. These results suggest that a DNA sequence-dependent means impact *RHO* expression, and that a precise base DNA sequence content and length (22 in humans and 23 bp in mice) ensures adequate quantitative gene expression levels of human and mouse RHO. It follows that the DNA base content and length linking CRX and NRL pairs generate a new code. Considering that the size of the DNA sequence space grows exponentially with sequence length (4^l^ for DNA, where l is the sequence length), the possible combinations are exceptionally high. However, the human and murine phylogenetic analysis has shown that they share in their respective DNA-linkers both sparse bases and almost complete conservation of the length of the DNA-linkers (Fig. 3 and S3). These aspects suggest that the DNA-linker code is evolutionarily constrained, and that evolution may have created the respective DNA-linkers after separation from the common ancestor to meet physiologically ranges of *RHO* expression relevant for the two species. To understand the possibilities and limitations of this new code, and to predict and control its activity (*55, 56*), it will be important to extend the analysis of the DNA-linkers exchanges between different species in a systematic way. Furthermore, the identification of genetic variants within DNA-linker compared with immediately adjacent sequences will be crucial to confirm its role in health and disease.

We additionally showed that DNA-linker-derived variation is interconnected with the binding sites of CRX and NRL. Individual mutation of the BSs of CRX and NRL abolishes the activity of the *RHO* promoter (Fig. 1B). In addition, changing their order and orientation led to a profound change in reporter expression activity (Fig. 2A). Although the BSs of CRX and NRL (as well as the DNA-linker itself) are not palindromic and can have four orientations relative to each other, they show complete conservation of both the sequence of the BSs and their orientation, supporting that these features are evolutionally constrained. Similarly, although linker DNA sequence conservation in mammals is poor, it is nevertheless evident that the DNA-linker orientation is also constrained. In fact, changing its orientation with respect to the CRX and NRL BSs, by reversing the 5’ - 3’ axis (Prom-L), greatly reduced the expression of the *RHO* reporter. However, this configuration modifies the 3’ and 5’ flanks of CRX and NRL, respectively, which, in turn may affect CRX and NRL binding. Indeed, the sequences flanking TFBSs are important for cooperative binding of TFs (*15, 16, 21*). Nonetheless, (i), the binding preferences of TF pairs are typically limited to a few bases flanking the TFBSs; (ii), the 3’ and 5’ flanks of the DNA-linker are variable along the mammalian phylogeny (Fig. 3A), and (iii), we observed that changes along the entire 22 bp length of the DNA-linker induce a wide variation in reporter gene expression. Thus, these aspects seem to be too broad both qualitatively and quantitatively to justify that what is observed here might be determined by the binding preferences of TF pairs alone, even considering mechanisms concerning the mutual influence of TF pairs that are exerted at greater distance (*66*). As opposed to short-mid range binding effects, the DNA-linker length appears too short to allow loop formation between CRX and NRL. Thus, we favor a model in which the DNA-linker is coupled in a continuum to its flanking CRX and NRL BSs. Nevertheless, the net contribution of binding on CRX and NRL TFBSs and the DNA-linker needs to be determined.

For there to be a mechanism of gene expression due to a contribution of the DNA sequence contained between two TFs, it must be possible to admit (and observe) that exist an independent and additional component in the DNA-protein interaction, distinct from the binding affinity of TFs for a DNA sequence, which is the only recognized process by which gene expression originates and occurs. The ability of both ZF-DB and KLF15 to block *RHO* expression *in vivo* at 20-fold lower concentrations than CRX and NRL (*55, 56*) and the unprotection from DNase of the DNA-linker sequence (*59, 60*) open the possibility that the DNA-linker, being not bound, may act subsequently of TF binding, and thus, by a DNA binding affinity-independent mechanism. Thus, in this view, our model admits that to the initial TFs binding of CRX and NRL occurring in affinity-dependent manner, a second event intervenes consequently (post-binding) as a DNA sequence-dependent intramolecular mechanism, which act as a third component adding to the two TFs activity.

The biochemical nature of this new DNA-TF interaction entanglement needs to be determined, including the dynamic regulation of the TF binding events, and relevant aspects related to the energy involved. These properties will also be instrumental to capture the basis of the interference produced by orthogonal DNA-binding proteins (ZF-DB-5 and KLF15). We previously have shown that ZF-DB-5 and KLF15 by binding the DNA-linker sequence block *RHO* expression regardless the side of the double strand bound (ZF-DB-5 minus strand, KLF15 plus strand, Fig. 4B) and the presence or absence of an effector domain (*61*). In a complementary manner, we have shown that reporter expression resulting from the same DNA-linker sequence arranged in the opposite direction (orientation 3’-5’, Prom-L) is severely impaired and yet, when bound by the same proteins ZF-DB-5 and KLF15, results in expression activation. Strikingly, this activation is more pronounced when the DNA-linker is bound by ZF-DB-5, which lacks catalytic domains. Thus, rather than a protein-protein steric interference these features may further support that ZF-DB-5 and KLF15 modify an activity passing through an exact DNA bases composition and length coupled to the action of two TFs.

Thus, beyond pattern and timing, we propose that levels of gene expression are contributed by DNA itself via a binding-independent mechanism which adds to the TFs binding-dependent mechanisms (Fig. S4). Once TFs are bound to TFBSs, by the specific DNA base sequence interposed between them allows the activity of TF-TFs to be integrated; as a result, the DNA base sequence and length constitute a new code that operates within a TF-DNA-TF complex, which thus represents a unit of activity, determining the appropriate final levels of gene expression. In this view, the role of DNA is thus threefold: it contains information that encodes for proteins, regulates their interactions (with a first feedback, binding to DNA), and participates in their own end action (with a second feedback, activity) by becoming a phenotype.

The control of gene expression proposed here may have evolved, and function more generally, as a still-hidden property, adding to the DNA-binding-dependent mechanisms determinant of gene expression.

## Acknowledgments

We thank Fondazione Telethon and the Telethon Institute of Genetics and Medicine (TIGEM) and the TIGEM Vector Core for vector production. We thank Irene Cantone and Pasqualina Colella for helpful discussions.

## Funding

This work was supported by the European Research Council/ERC grant 311682 “Allelechoker” (to EMS); ERC proof-of concept (ERC-POC). “Insight” 813223 (to EMS); Missione 4 - Center for Gene Therapy and Drugs based on RNA Technology-WP3.5 Stroke and other brain diseases, WP 3.5.7 (to EMS); Prin MUR: 2022LBZSTR (to EMS).

## Author contributions

EMS conceived the study. RM, MP, SB, EMS, designed the experiments and analyzed data. MS, EM, performed histological analysis. GD performed bioinformatic analysis. EM, EMS, MLB performed in vivo studies. RM, AM, ED, CM and EMS draft the manuscript. RM, AM, ED, MP, SB, EM, MR, performed the molecular biology experiments. EMS wrote the manuscript and supervised the study.

## Competing interests

E.M.S., S.B. and E.M. are inventors on patents, “Artificial DNA-binding proteins and uses thereof”, WO2015075154A3, US20160289284A1. E.M.S., S.B. and E.M. are inventors on a pending patent “Ectopically expressed transcription factors and uses thereof”, PCT/EP2018/086782, Department of Translational Medical Sciences (DiSMeT) University of Naples “Federico II”, Italy, which may encompass the findings. The remaining authors declare no competing interests.

## Data and materials availability

All data are available in the main text or the supplementary materials.

## Notes

### Competing Interest Statement

The authors have declared no competing interest.

### Summary of Updates

This version of the manuscript contains updates on the subject.

