## Supplementary for "A DNA base-specific sequence interposed between CRX and NRL contributes to *RHODOPSIN* expression"

### Materials and Methods

#### Plasmid construction

The human rhodopsin proximal promoters (hRHOp, 164 bp from the transcription starting site (TSS) and 95bp of 5'UTR) and the mouse proximal promoter (mRHO, 164bp from the transcriptional starting site (TSS) and 78bp of 5'UTR) were generated by gene synthesis of Eurofins MWG® and cloned in pAAV2.1-eGFP using NheI and NotI restriction enzymes. All variant promoters of hRHOp (Prom-3A, Prom-H, Prom-Me, Del-3', Del-C, Prom-E, Prom-Q, Prom-N, Del-5', Prom-S, hRHOp-mRRU, mRHOp-hRRU, PromG-1, PromG-2, Prom-L, MutCRX1, MutCRX2, MutCRX1.2, MutNRL) were generated by gene synthesis of Eurofins MWG® and cloned in pAAV2.1-eGFP using NheI and NotI restriction enzymes.

#### AAV vector preparation and purification

AAV vectors were produced by by the TIGEM AAV Vector Core by triple transfection of HEK293 cells followed by two rounds of CsCl2 purification. For each viral preparation, physical titers [genome copies (GC)/mL] were determined by averaging the titer achieved by dot-blot analysis and by PCR quantification using TaqMan (Applied Biosystems, Carlsbad, CA, USA) (*68*).

#### Mice

All procedures were performed in accordance with institutional guidelines for animal research and all of the animal studies were approved by the authors. C57BL/6 mice (Charles Rivers Laboratories, Calco, Italy) and P347S+/+ animals (REF. Li T, Snyder WK, Olsson JE, Dryja TP. Transgenic mice carrying the dominant rhodopsin mutation P347S: evidence for defective vectorial transport of rhodopsin to the outer segments. *Proc Natl Acad Sci USA*. 1996;93(24):14176–14181.) were bred in the animal facility of the Biotechnology Centre of the Cardarelli Hospital (Naples, Italy). P347S–/– mice were crossed with C57BL6/J mice (Charles Rivers Laboratories) to obtain the P347S+/– mice (*55*).

**Vector administration**
Intraperitoneal injection of ketamine and medetomidine (100mg/kg and 0.25mg/kg respectively), then AAV vectors were delivered sub-retinal via a trans-scleral transchoroidal approach. (*55*).
Each eye was co-injected with two AAV vectors: AAV vector carrying AAV-eGFP driven by human and murine proximal promoter variants (dose $1x10^9$GC/µl) and an AAV control vector encoding for human *RHO* gene driven by the GNAT1 proximal promoter, or AAV control vector encoding for mCherry driven by hGNAT1 proximal promoter (dose $1x10^8$GC/ µl) (*55*). Mouse experimental groups were sized at $n \geq 4$. P347S mice subretinally injected at post-natal day 14 (PD14) with AAV8-CMV-ZF-DB, AAV8-CMV-ZF-DB-5, AAV8-CMV-KLF15, or AAV8-CMV-eGFP and analysed at P30 as described (*55*, *56*).

**qReal Time PCR**
RNAs from tissues were isolated using RNAeasy Mini Kit (Qiagen), according to the manufacturer protocol. cDNA was amplified from 1 µg isolated RNA using QuantiTect Reverse Transcription Kit (Qiagen), as indicated in the manufacturer instructions. The PCRs with cDNA were carried out in a total volume of 20 µl, using 10 µl LightCycler 480 SYBR Green I Master Mix (Roche) and 400nM primers under the following conditions: pre-Incubation, 50°C for 5 min, cycling: 45 cycles of 95°C for 10 s, 60°C for 20 s and 72°C for 20 s. Each sample was analyzed in duplicate in two-independent experiments. Transcript levels of *eGFP* and h*RHO* were normalized against murine Gapdh and Actβ (DCt), and then the eGFP expression shown is obtained as fold change over control (co-injected vector). Furthermore, the AAV-hGNAT1-hRHO was co-injected to normalize the variability due to subretinal administration. We used the following primers: Act_forward (CAAGATCATTGCTCCTCCTGA) and Act_reverse (CATCGTACTCCTGCTTGCTGA), Gapdh_forward (GTCGGTGTGAACGGATTTG) and Gapdh_reverse (CAATGAAGGGGTCGTTGATG); eGFP_forward (ACGTAAACGGCCACAAGTTC) and eGFP_reverse (AAGTCGTGCTGCTTCATGTG); hRHO_forward (TCATGGTCCTAGGTGGCTTC) and hRHO_reverse (GGAAGTTGCTCATGGGCTTA); mCherry_forward (CACTACGACGCTGAGGTCAA) and mCherry_reverse (GTGGGAGGTGATGTCCAACT).

**Histological analysis**
For morphological studies, after the mice were killed by cervical dislocation, the eyecups were harvested, fixed by immersion in 4% paraformaldehyde, and then embedded in OCT (KalteK). For each eye, 150 to 200 serial sections (10-µm thick) were cut along the horizontal plane; the sections were progressively distributed on 10 glass slides so that each slide contained 15 to 20 sections representative of the whole eye at different levels. Slides were coverslipped with Vectashield containing DAPI (4',6-diamidino-2-phenylindole; Vector laboratories, Burlingame, CA, USA) to stain cells nuclei and retinal histology was analyzed a Leica Fluorescence Microscope System (Leica Microsystems GmbH, Wetzlar, Germany).

**Transient transfection and analysis of promoter activity by qReal Time PCR**
HEK293 cells were plated in six-well plates at a density of $1x10^6$ cells/well in Dulbecco’s modified Eagle’s medium supplemented with 10% fetal bovine serum and 1% penicillin/streptomycin and grown for 24h a 37° C. The cells were transfected at 80% confluency using a TransIT-X2 Dynamic Delivery System (MIRUS), as indicated in the manufacturer instructions. We co-transfected the following

constructs: 50 ng reported plasmid, 250ng of a plasmid encoding for human CRX, 125 ng of a plasmid expressing human NRL and 125ng CMV-ZF5 or CMV-KLF15 plasmid. The DNA amount was kept at 550 ng/well with pUC19. Cells were grown and harvested 48h post transfection in TRIzol™ Reagent (Life Technologies). The cDNA was amplified from 1 µg isolated RNA using QuantiTect Reverse Transcription Kit (Qiagen), as indicated in the manufacturer instructions. The PCRs with cDNA were carried out in a total volume of 20 µl, using 10 µl LightCycler 480 SYBR Green I Master Mix (Roche), 400nM primers under the following conditions: pre-Incubation, 50°C for 5 min, cycling: 45 cycles of 95°C for 10 s, 60°C for 20 s and 72°C for 20 s. Each sample was analysed in triplicate. Transcript levels of cells were measured by real-time PCR using the LightCycler 480 (Roche) and the following primers: PCR eGFP_forward (ACGTAAACGGCCACAAGTTC) and eGFP_reverse (AAGTCGTGCTGCTTCATGTG), GAPDH_forward (GAAGGTGAAGGTCGGAGT) and GAPDH_reverse (GAAGATGGTGATGGGATTTC). The eGFP expression shown is obtained normalizing to GAPDH.

**Statistical analyses**. Data are presented as mean ± Error bars indicate standard error mean (SEM). Statistical significance was computed using the One-Way Anova test (p-values ≤ 0.05), and post-hoc Dunnett's multiple comparisons test (p-values ≤ 0.05). Only in figure 2B we performed t-student test (p-values ≤ 0.0001).

**Motif Scan**

TF human motifs were retrieved from HOCOMOCO (version 11) (*63*) and scanned on target sequences using FIMO (meme-suite 5.5) (Grant CE,2011), with background model derived from target sequence composition and parameters --norc and –thresh 0.01.

Matches with positive scores and p-value<0.01 were considered. Alternative matches for a given TF on the same sequence were disambiguated by considering the match with the highest score.

**TF expression in retina**

Transcript and protein expression data were retrieved from the Human Protein Atlas version 23.0 (https://www.proteinatlas.org/). We considered as positively expressed transcripts with normalized TPM >1 in human retina (n=13.114), and proteins with positive detection in photoreceptor cells (n=79).

**Electrophysiological testing**

The method is as described (*67*). Briefly, mice were dark reared for 3 hr and anesthetized. Flash electroretinograms (ERGs) were evoked by light flashes generated through a Ganzfeld stimulator (CSO, Costruzione Strumenti Oftalmici, Florence, Italy). ERG analysis in scotopic conditions were evoked by 11 stimuli (from -4 to +1.3 log cd*s/$m^2$) with an interval of 0.6 log unit. and registered as previously described (*55*).

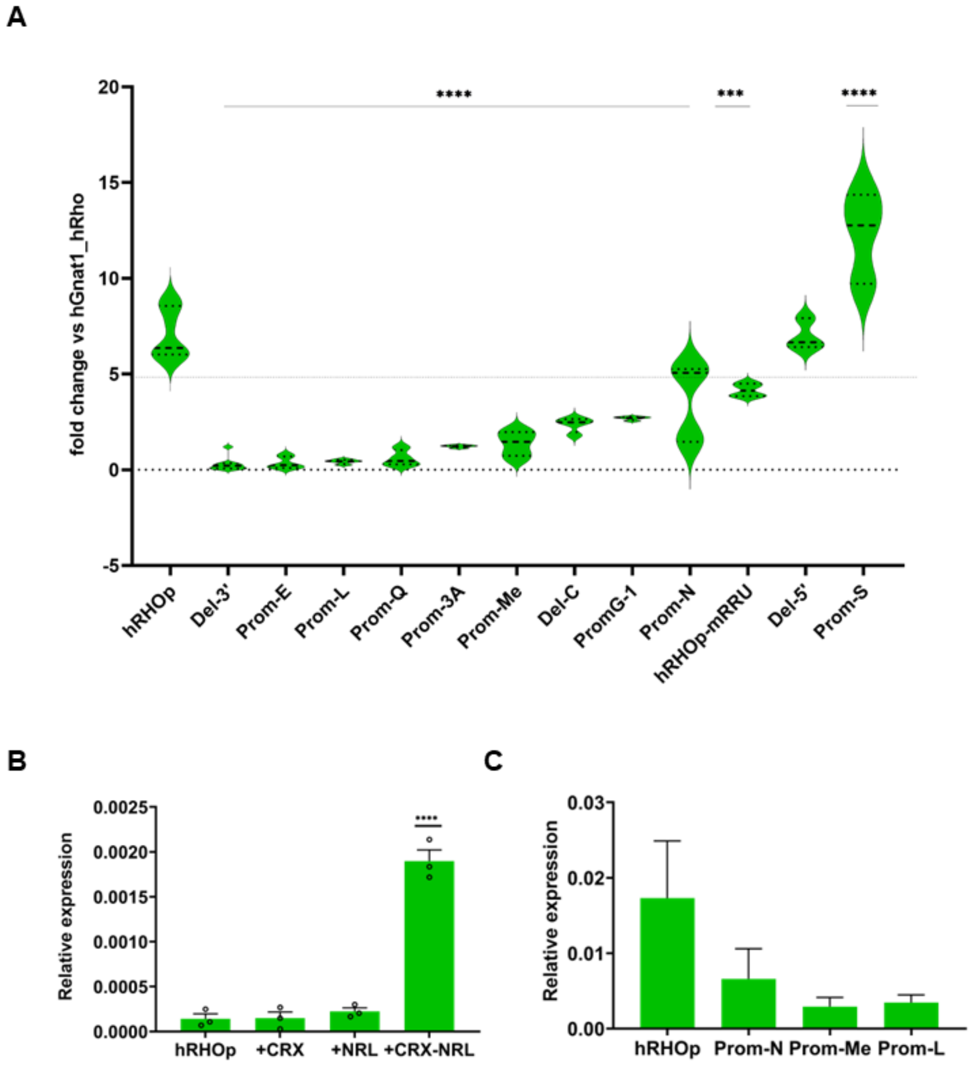


**Fig. S1.**

**(A)** Cumulative *eGPF* variations across the promoters tested in vivo, normalized to wild-type promoter. **(B)** To test promoter activation in vitro the cells HEK-293 were transfected with hRHOp and its transactivating transcription factors CRX and NRL. **(C)** In vitro (HEK-293) transfection of hRHOp and mutants as indicated.

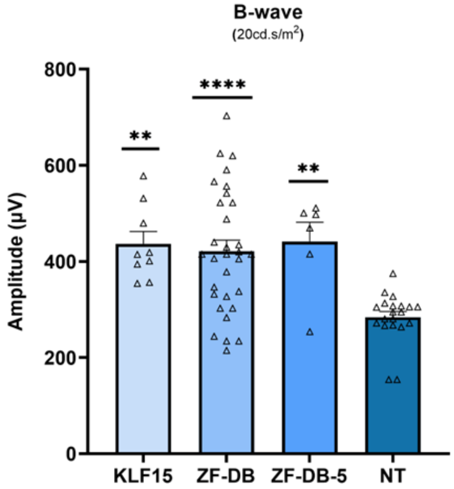


**Fig. S2.**

Electroretinography (ERG) analysis on P347S mice subretinally injected at post-natal day 14 (PD14) with AAV8-CMV-ZF-DB (n=10), AAV8-CMV-ZF-DB-5 (n=6), AAV8-CMV-KLF15 (n=9), or AAV8-CMV-eGFP (n=20) and analysed at P30. Preservation of retinal maximal responses, scotopic (dim light) B-waves amplitudes evoked by light flash at +1.3 log cd*s/$m^2$ (which correspond to 1x10-5.2 to 20.0 cd*s/$m^2$), were preserved in AAV8-CMV-ZF-DB, ZF-DB-5 and KLF15 compared to eGFP control.

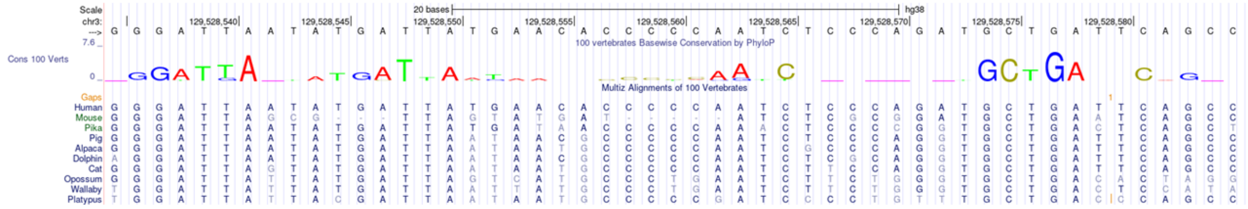


**Fig. S3.**

**Evolutive conservation of *RHO* promoter.** Upper panel: logo of basewise conservation by PhyloP. Lower panel: multiple sequence alignment of representative mammalian species among 100 vertebrates.

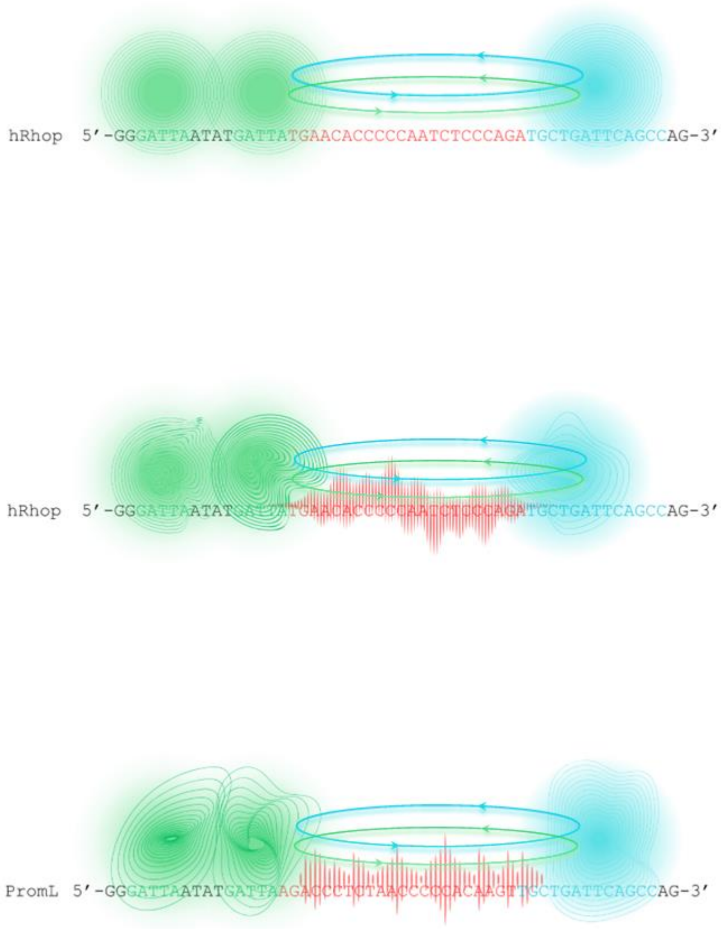


**Fig. S4.**

**Graphical representation modelling the novel DNA-TF interaction mechanism proposed.** Top drawing shows standard model (TFs binding-dependent mechanisms) of TFs interaction with their respective binding sites. CRX and NRL TFs and their binding sites are represented in green and light blue respectively, and their TF-TF interaction is represented by light blue and green arrows. In the middle and bottom drawings, the binding-independent interaction mechanism proposed, mediated by the DNA itself. Distinct DNA bases composition (and length) of the DNA-linker (red sequence; middle drawing, hRHOp DNA-linker; bottom, hRHOp DNA-linker arranged in the opposite orientation 5'-3', Prom-L in the text) differentially impact (red waves) on TF-TF activity adding to TFs binding-dependent mechanisms (top drawing), eventually leading to differential gene expression levels.
